# Gz Enhanced Signal Transduction assaY (G_Z_ESTY) for GPCR deorphanization

**DOI:** 10.1101/2024.07.26.605282

**Authors:** Luca Franchini, Joseph J. Porter, John D. Lueck, Cesare Orlandi

## Abstract

G protein-coupled receptors (GPCRs) are key pharmacological targets, yet many remain underutilized due to unknown activation mechanisms and ligands. Orphan GPCRs, lacking identified natural ligands, are a high priority for research, as identifying their ligands will aid in understanding their functions and potential as drug targets. Most GPCRs, including orphans, couple to G_i/o/z_ family members, however current assays to detect their activation are limited, hindering ligand identification efforts.

We introduce G_Z_ESTY, a highly sensitive, cell-based assay developed in an easily deliverable format designed to study the pharmacology of G_i/o/z_-coupled GPCRs and assist in deorphanization. We optimized assay conditions and developed an all-in-one vector employing novel cloning methods to ensure the correct expression ratio of G_Z_ESTY components. G_Z_ESTY successfully assessed activation of a library of ligand-activated GPCRs, detecting both full and partial agonism, as well as responses from endogenous GPCRs. Notably, with G_Z_ESTY we established the presence of endogenous ligands for GPR176 and GPR37 in brain extracts, validating its use in deorphanization efforts. This assay enhances the ability to find ligands for orphan GPCRs, expanding the toolkit for GPCR pharmacologists.

## 1. INTRODUCTION

In mammals, G Protein Coupled Receptors (GPCRs) are the largest family of cell surface receptors and are involved in a wide range of physiological and pathological processes^1,2^. Orphan GPCRs are those whose natural ligands have not yet been identified or agreed upon, making them difficult to study. Deorphanization, the process of identifying endogenous ligands for orphan GPCRs, is often achieved by analyzing receptor activation by candidate ligands in heterologous cell systems. These candidate ligands are selected based on their overlapping *in vivo* localization with a target GPCR, by isolation from tissue extracts that exert measurable physiological effects, or via screening of large libraries of endogenous compounds including small molecules, peptides, and lipids with a demonstrated biological activity^3–11^. Research that led to the deorphanization of the Glucagon-Like Peptide-1 (GLP1) receptor illustrates how identifying a GPCR’s endogenous ligand can unlock new therapeutic opportunities. Initially discovered as a gut-derived hormone that enhances insulin secretion in response to elevated plasma glucose levels^12^, GLP1 was found to bind to a putative receptor in the brain and pancreas^13,14^. This discovery led to the cloning of its receptor, a GPCR, in the early 1990s^15–17^. Since then, numerous GLP1 receptor agonists have been approved for treating type 2 diabetes mellitus and, more recently, for chronic weight management in adults with obesity^18–20^.

A central challenge in the process of deorphanization is the choice of a measurable outcome of receptor activation. Cell-based assays exploiting various signaling properties and readouts have been developed, each with their own advantages and limitations. These assays measure proximal events such as heterotrimeric G protein dissociation^21–24^ or β-arrestin recruitment^25,26^; they quantify the accumulation of second messengers^27–30^; or they utilize genetic reporters that depend on transcriptional regulation events initiated by GPCRs^31,32^. Further sets of assays are label-free and agnostic to the molecular mechanism leading to the change being measured^33,34^. All of these assays differ in sensitivity and time required to obtain a measurable signal. Most importantly, when studying orphan GPCRs, lack of information about G protein coupling profile or ability to recruit β-arrestins requires further considerations. To partially address this issue, we recently quantified the constitutive activity of several orphan GPCRs and showed that many are coupled to G proteins belonging to the G_i/o/z_ family^35^.

Here, we performed an in-depth optimization of a cell-based assay aimed at measuring G_i/o/z_ coupled receptor activation. We demonstrated an improved sensitivity and the wide applicability of this assay that we named G_z_ Enhanced Signal Transduction assaY (G_z_ESTY). We generated and tested an all-in-one plasmid for the mammalian expression of each component required for G_z_ESTY at the optimized ratio. Finally, we applied this assay to demonstrate the presence of endogenous ligands for orphan receptors GPR176 and GPR37 in unfractionated mouse brain extract.

## 2. METHODS

### 2.1 Cell cultures and transfections

HEK293T/17 cells were purchased from American Type Culture Collection (ATCC) and cultured at 37°C and 5% CO_2_ in Dulbecco’s Modified Eagle’s Medium (DMEM; Gibco, 10567-014) supplemented with 10% fetal bovine serum (FBS; Biowest, S1520), Minimum Eagle’s Medium (MEM) non-essential amino acids (Gibco, 11140-050), sodium pyruvate (Gibco, 11360-070), GlutaMAX (Gibco, 35050-061), and antibiotics (100 units/ml penicillin and 100 µg/ml streptomycin; Gibco, 15140-122). 1 × 10^6^ cells/well were seeded in 6-well plates in medium without antibiotics. After 4 hours, cells were transfected using linear 25 kDa polyethylenimine (PEI) (Polysciences; 23966) at a 1:3 ratio between total µg of DNA plasmid (2.5 µg) and µl of PEI (7.5 µl). A pcDNA3.1 empty vector was used to normalize the amount of transfected DNA. Charcoal-stripped FBS (S162C) and dialyzed FBS (347G18-D) were purchased from Biowest.

### 2.2 DNA constructs and cloning

Plasmids encoding Dopamine D2 Receptor (D2R), GABAB receptor subunits (GABBR1 and GABBR2), α2-adrenergic receptor 2A (ADRA2A), µ-opioid receptor (MOR), δ-opioid receptor (DOR), κ-opioid receptor (KOR) α2-adrenergic receptor 2B (ADRA2B), α2-adrenergic receptor 2C (ADRA2C), serotonin receptor 1B (5-HT1B), masGRK3CT-Nluc, PTX-S1, and Gα_oA_ were generous gifts from Dr. Kirill Martemyanov (UF Scripps Institute, FL). Gβ_1_-Venus^156–239^ and Gγ_2_-Venus^1-155^ were generous gifts from Dr. Nevin Lambert (Augusta University, GA) ^22^. Codon-optimized coding sequences of the following receptors were a kind gift from Dr. Bryan Roth (University of North Carolina, NC; Addgene kit no. 1000000068) ^31^ and were used for subcloning into pcDNA3.1 plasmids removing V2-tail, TEV site, and tTA sequences: follicle-stimulating hormone receptor (FSHR), cannabinoid receptor 1 (CB1R), cannabinoid receptor 2 (CB2R), sphingosine-1 phosphate receptor 1 (S1PR1), lysophosphatidic acid 2 receptor (LPAR2), somatostatin receptor 1 (SSTR1), neuropeptide FF receptor 1 (NPFFR1), free fatty acid receptor 3 (FFA3R), galanin 1 receptor (GALR1), formyl peptide receptor 1 (FPR1), hydroxycarboxylic acid receptor 2 (HCAR2), neuropeptide Y receptor Y1 (NPYR1), protease-activated receptor 1 (PAR1), protease-activated receptor 2 (PAR2), histamine receptor H3 (HRH3), dopamine D1 receptor (D1R), dopamine D3 receptor (D3R), adenosine A2B receptor (ADORA2B), parathyroid hormone 1 receptor (PTH1R), melanocortin 4 receptor (MC4R), and calcitonin-like receptor (CLR). Plasmids encoding the cAMP sensor (pGloSensor™-22F) and CRE-Nluc reporter were purchased from Promega. pFL-tk encoding firefly luciferase under the control of constitutively active thymidine kinase promoter was used as a normalizer for reporter assays. The plasmids encoding the following G_s_-derived chimeras were a kind gift from Dr. Robert Lucas (University of Manchester, UK)^27^: GsGz (Addgene plasmid #109355), GsGo (Cys) (Addgene plasmid #109375), GsGi (Cys) (Addgene plasmid #109373). pOZITX-S1 was a kind gift from Jonathan Javitch (Addgene plasmid #184925). All constructs were verified by Sanger sequencing.

A modular plasmid system was used to assemble the components of G_Z_ESTY into single multicomponent plasmids. Five insert shuttle plasmids (P1-P5) and a destination plasmid were constructed with matching homology regions for Gibson assembly flanking an insert sequence. For assembly, the shuttle plasmids and destination plasmid were digested with the Type IIS restriction enzyme CspCI, which possesses a relatively rare 7-bp recognition sequence. As CspCI is a type IIS restriction sequence, it cleaves outside of its recognition sequence, allowing for generation of DNA fragments containing unique ends for homology-directed assembly (i.e. Gibson assembly). P1 contained a short CMV promoter (lacks CspCI site found in conventional CMV promoter) driving expression of GloSensor-22F, P2 contained a short ubiquitin C promoter (UbC) driving expression of PTX-S1, P3 contained an ampicillin selection marker and a short CMV promoter driving expression of the receptor of interest, P4 was generally a blank insert except for the GABAB receptor for which P4 contained a short CMV promoter driving expression of the GABAB receptor subunit 2 (with P3 containing subunit 1), and P5 contained a UbC promoter driving expression of the GsGz chimera. To generate plasmids only containing some of these inserts of interest, blank inserts were used for any of P1-P5 containing G_Z_ESTY components not required. The destination plasmid was a pUC57-Kan containing the homologous sequences for Gibson assembly of the inserts. Of note, a plasmid with an alternate resistance marker to AmpR was used as this allows for selection of the final assembled clone without any carry-through of undigested insert plasmids. The CspCI digest contained 110 fmol of each insert plasmid (∼375 ng for a 5 kb insert plasmid), 37.5 fmol of the destination plasmid (∼125 ng for a 5 kb destination plasmid), 5 µL rCutSmart buffer (NEB), and 1 µL CspCI (5,000 units/mL; NEB) in a 50 µL reaction, incubated at 37 °C for 1 hour, and heat inactivated at 65 °C for 20 minutes. For the Gibson assembly, 5 µL of the CspCI digest was mixed with 5 µL of NEBuilder HiFi DNA Assembly Master Mix (NEB), incubated at 50 °C for 15 minutes. For transformation of the assembled plasmid, 1.5 µL of the Gibson assembly mix was transformed into either NEB 5-alpha Competent *E. coli* (NEB) or NEB 10-beta Electrocompetent *E. coli* (NEB) and plate on selective LB agar containing both kanamycin (selects for destination plasmid backbone) and ampicillin (selects for P3 insert).

### 2.3 Chemicals

The following chemicals were purchased: clonidine (Tocris), dopamine (Tocris), GABA (Tocris), serotonin (Tocris), 1-oleoyl lysophosphatidic acid (Tocris), DAMGO (MedChemExpress), TFLLR (MedChemExpress), human PAR-2 (1-6, SLIGKV) (MedChemExpress), human galanin (1-30) (MedChemExpress), IBMX (MedChemExpress), SEW2871 (MedChemExpress), somatostatin-14 (Cpc Scientific), human neuropeptide Y (13-36) (Cpc Scientific), neuropeptide FF (Thermo Scientific Chemicals), MK-6892 (MedChemExpress), 2-arachidonoyl glycerol (Cayman Chemicals), N-Formyl-Met-Leu-Phe (R&D systems), SNC80 (Adipogen), isobutyric acid (TCI chemicals), salvinorin A (ChromaDex Inc.), morphine (Mallinckrodt Chemical Company), human β-endorphin (Sigma-Aldrich), teriparatide (MedChemExpress), NDP-α-MSH (Phoenix Pharmaceuticals), AB-MECA (MedChemExpress), quinpirole (Tocris), calcitonin gene-related peptide (CGRP) (AnaSpec).

### 2.5 cAMP measurements using G_Z_ESTY

HEK293T/17 cells were transfected as described above with pGloSensor™-22F cAMP plasmid (Promega) and indicated plasmids for mammalian expression of GPCRs, PTX-S1 (or OZITX-S1), and Gs-based chimeras (GsGo, GsGi, or GsGz). 18 hours post-transfection, cells were mechanically detached using a gentle stream of PBS, centrifuged at 500 g for 5 minutes, and resuspended in 300 µl of PBS containing 0.5 mM MgCl_2_ and 0.1% glucose. 40 µl of the cell suspension were transferred to each well of 96-well flat-bottomed white plates (Greiner Bio-One). D-Luciferin potassium salt (Gold Bio; #DLUCK100) was dissolved in HEPES buffer 10 mM pH 7.4 to obtain a 5x stock at 30 mg/mL. 10 µL of 5X D-luciferin solution were added to the cells. Cells were then incubated at 37°C and 5% CO_2_ for 1 hour and then equilibrated in the plate reader at 28°C, until stable baseline values were reached (∼15 minutes). The luminescence signal was monitored approximately every 30 seconds using a POLARstar Omega microplate reader (BMG Labtech).

In assays performed on coated 96 well plates, cells were transfected in 6-well plates. The day after transfection, the media was removed, and cells were gently washed with PBS once and briefly trypsinized with 100 µL of trypsin. Trypsinization was blocked by the addition of 900 µL of transfection media containing 10% FBS, cells were counted and seeded at 6.5 × 10^4^ cells/well on a sterile white 96-well plate coated with poly-D-lysine. Cells were then incubated overnight at 37°C and 5% CO_2_. 24 hours later, media in each well was replaced with 40 µL of PBS containing 0.5 mM MgCl_2_, 0.1% glucose, and 10 µL of 5X D-Luciferin, and then incubated for 1h at 37°C. The 96-well plate was moved into the plate reader, equilibrated at 28°C until stable luminescence values were obtained and finally treated according to the experiment setup.

### 2.6 G protein nanoBRET assay

GPCR activation in live cells was measured as BRET signal between Venus-Gβ1γ2 and masGRK3CT-Nluc performed as described previously^23^. 2.5 µg of total DNA was transfected according to the following ratio: 0.21 µg of Gβ1-Venus(156-239); 0.21 µg of Gγ2-Venus(1-155); 0.21 µg of masGRK3CT-Nluc; 0.42 µg of Gα_oA_ proteins; and 0.21 µg of receptor. Empty vector pcDNA3.1 was used to normalize the amount of transfected DNA. 18 hours after transfection, HEK293T/17 cells were washed once with PBS. Cells were then mechanically harvested using a gentle stream of PBS, centrifuged at 500 g for 5 minutes, and resuspended in 500 µl of PBS containing 0.5 mM MgCl_2_ and 0.1% glucose. 25 µl of resuspend cells were distributed in 96-well flat-bottomed white microplates (Greiner Bio-One). The Nluc substrate, furimazine (N1120; Promega) was used according to the manufacturer’s instructions. Luminescence was quantified at room temperature using a POLARstar Omega microplate reader (BMG Labtech). BRET signal was determined by calculating the ratio of the light emitted by Venus-Gβ1γ2 (collected using the emission filter 535/30) to the light emitted by masGRK3CT-Nluc (475/30). The average baseline value (basal BRET ratio) was recorded for 5 seconds before agonist application.

### 2.7 CRE luciferase reporter assay

HEK293T/17 cells were plated at a density of 1 × 10^6^ cells/well in 6-well plates in an antibiotic-free medium and transfected as described above. 2.5 µg of total DNA was transfected according to the following ratio: 0.10 µg of pFL-tk plasmid expressing firefly luciferase under control of the constitutive thymidine kinase promoter; 0.21 µg of CRE-Nluc luciferase reporter; 1.98 µg of GPCR; and only in experiments screening G_i/o_ activation, 0.21 µg of GsGi1 chimera. pcDNA3.1 was used to normalize the amount of transfected DNA. Cells were incubated overnight and then serum-starved in Opti-MEM for 4 hours before collection. Transfected cells were harvested using a gentle stream of PBS, centrifuged for 5 minutes at 500 ×g, and resuspended in 150 µl of PBS containing 0.5 mM MgCl_2_ and 0.1% glucose. 30 µl of cells were incubated in 96-well flat-bottomed white microplates (Greiner Bio-One) with 30 µl of luciferase substrate according to manufacturers’ instructions: furimazine (Promega NanoGlo; N1120) for Nluc, and D-luciferin (Promega BrightGlo; E2610) for firefly luciferase. Luciferase levels were quantified using a POLARstar Omega microplate reader (BMG Labtech). Firefly luciferase expression was used to normalize the signal and compensate for variability due to transfection efficiency and number of cells.

### 2.8 Mice

C57BL/6J mice were housed in a temperature and humidity-controlled room in the vivarium of the University of Rochester on a 12:12-h light/dark cycle (lights off at 18:00 h) provided with food and water ad libitum. Male and female adult mice were used for the experiments. Mice were sacrificed by cervical dislocation, the head was removed and placed immediately in a microwave at maximum power for 10s to block protease activity^36^. One brain was removed and immediately homogenized in a glass-glass homogenizer with 4mL of PBS. Homogenate was then sonicated three times on ice for 10s at 30% power and spun for 10 min at 14,000g at 4°C. 50 µL of supernatant was then applied to the cells. All procedures were pre-approved and carried out following the University Committee on Animal Resources (UCAR) at the University of Rochester. Bovine pituitary extract was purchased from MP biologicals (cat#2850450).

### 2.9 Statistical analysis

Statistical analysis was performed using GraphPad Prism 9 software. The number of replicates and type of statistical analysis used are described in each figure legend.

## 3. RESULTS

### 3.1 G_s_-based protein chimeras combined with a cAMP biosensor allow for the measurement of the activation of G_i/o/z_-coupled receptors

Several cell-based assays are currently available to study the activation of G_i/o/z_-coupled receptors. In the G protein nanoBRET assay, receptor activation in response to ligands is measured as an increase in BRET signal due to the release of Gβγ-venus and its interaction with a membrane-anchored biosensor that consists of the C-terminus of GRK3 fused to Nluc (**Supplementary Figure 1A**)^22,35^. The main advantage of this assay is the rapid measurement of a proximal event in GPCR activation that allows for real-time analysis of the receptor kinetics but with a trade-off in assay sensitivity (**Supplementary Figure 1D**). Genetic reporters expressing luciferase under the control of a GPCR-inducible promoter are commonly used. These assays have been developed to detect signals downstream of GPCR coupling to G_s_ (CRE-luc reporter), G_q_ (NFAT-luc reporter), and G_12/13_ (SRE-luc reporter)^32^. Activation of G_i/o/z_-coupled receptors can be quantified as a reduction in the luciferase accumulation using a CRE-luc reporter, or indirectly as Gβγ modulation of signaling pathways regulating the SRE-luc reporter. A more sensitive approach adopts G protein chimeras based on the core of Gα_s_ or Gα_q_ and the last few amino acids of members of the G_i/o/z_ family^37,38^. This strategy allows the detection of an accumulation of luciferase using CRE-luc and NFAT-luc, respectively, in response to the activation of G_i/o/z_-coupled receptors (**Supplementary Figure 1B**). One limitation of this approach is that the receptor must be exposed to an agonist for an extended time to accumulate measurable quantities of luciferase in the cell (**Supplementary Figure 1D**). During this time, processes like receptor desensitization, internalization, and compensatory mechanisms can take place in the cell reducing the assay reliability. A good compromise between speed and assay sensitivity is based on measuring the accumulation of second messengers. Activation of G_i/o/z_-coupled receptors inhibits the activity of adenylyl cyclase thereby reducing cAMP accumulation in response to forskolin treatments. Real-time measurements of cAMP levels can rapidly indicate if a G_i/o/z_-coupled receptor has been activated. This assay depends on the side-by-side comparison of cAMP accumulation in cells treated with an agonist and in cells treated with vehicle. Such an approach is rapid, achieving a signal plateau within 15 minutes of agonist application (**Supplementary Figure 1C-D**). Alternatively, co-expression of Gα_s_-based chimeras bearing the C-terminus of G_i/o/z_ family members can be used to redirect receptor activation to stimulate adenylyl cyclase and cAMP accumulation (**Supplementary Figure 1C-D**)^27^. Overall, this last strategy yields an optimal balance between assay duration and sensitivity.

Building on these observations, we tested the activation of four prototypical G_i/o/z_-coupled GPCRs, dopamine D2 receptor (D2R), α2 adrenergic receptor (ADRA2A), γ-aminobutyric acid B receptor (GABAB), and µ opioid receptor (MOR) using three Gα_s_-based chimeras bearing the C-terminus of Gα_i_, Gα_o_, and Gα_z_. Application of selective agonists revealed a robust accumulation of intracellular cAMP as measured using a split-luciferase GloSensor (**Figure 1A**). We measured baseline luminescence for 3 minutes before applying each agonist as well as the maximal amplitude obtained in response to agonist stimulation (**Figure 1B**). Using these measurements, we calculated the fold change induction for each condition as an index that we can use to compare sensitivity between assays (**Figure 1C**). These experiments revealed that the GsGz chimera always showed a low baseline that contributed to the highest fold change observed with three out of four receptors.

**Figure 1.**
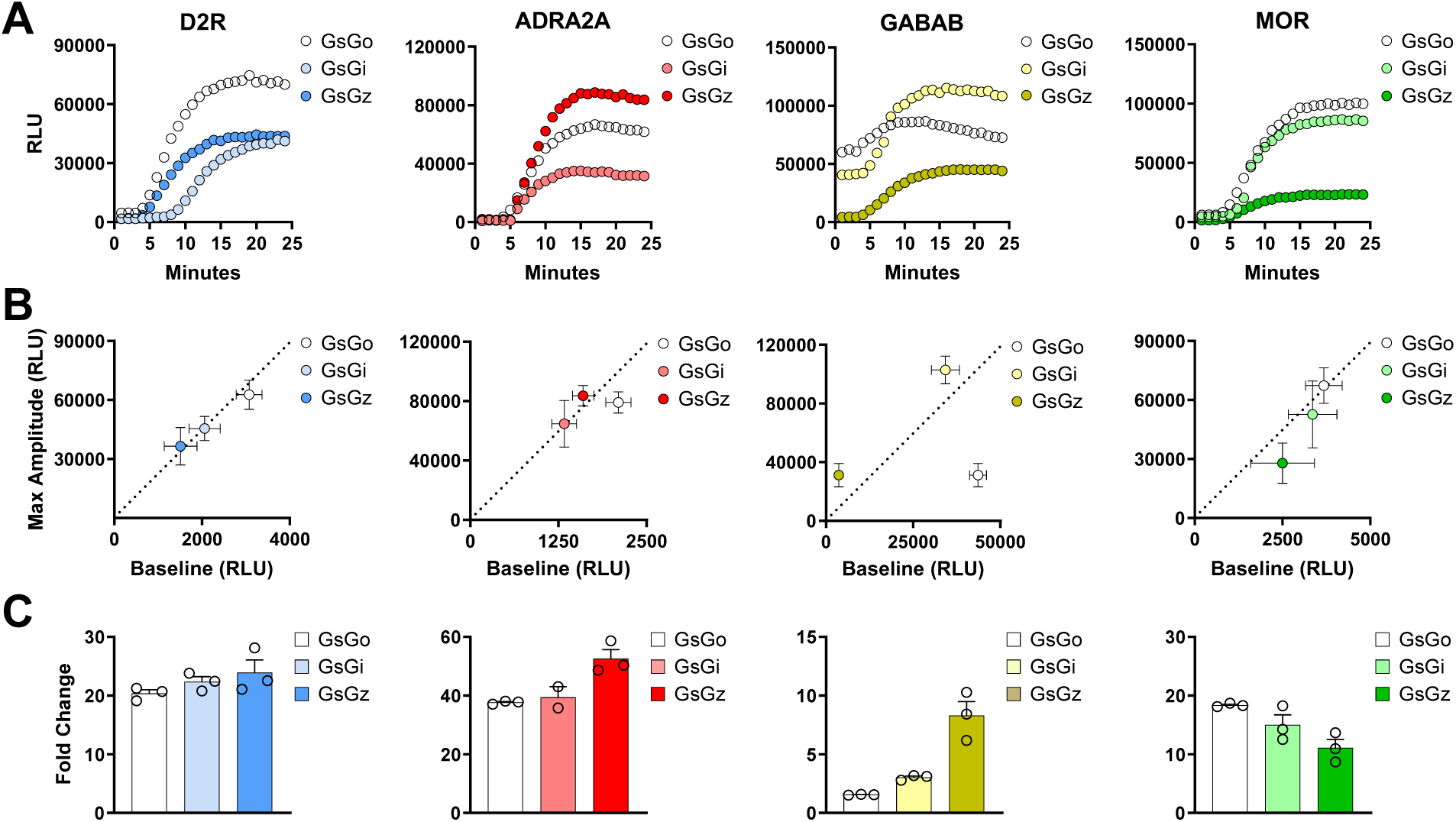
Real-time activation of Gs-based chimeras by G_i/o/z_-coupled receptors. (**A**) Representative kinetics of cAMP accumulation in response to agonist stimulation of indicated GPCRs using 10 µM dopamine, 10 µM clonidine, 10 µM GABA, and 1 µM DAMGO, respectively. (**B**) Correlation between maximal amplitude and basal signal for each receptor co-expressed with indicated G protein chimeras. (**C**) Fold change calculated as the ratio of maximal amplitude over baseline reported in panel **B**. Data are shown as means ± SEM. N=3 independent replicates.

### 3.2 The assay sensitivity shows a clear dependence on both the transfection ratio of assay components and the inhibition of endogenous G_i/o_ proteins

We optimized the assay conditions by adjusting the amount of transfected plasmids used to express each Gα_s_-chimera. (**Figure 2A**). We found that, generally, excessive amounts of Gα_s_-chimeras result in higher baselines, which attenuate the observed fold change (**Supplementary Figure 2**). Moreover, considering that GsGz, just like Gα_z_, is insensitive to the action of pertussis toxin (PTX), we introduced it in the assay aiming to reduce the adenylyl cyclase inhibition due to endogenous Gα_i/o_ proteins. We found that very low levels of transfected PTX boosted the induction of cAMP for each GPCR from a minimum of 1.5 times (GABABR) to a maximum of 6.1 times (D2R) (**Figure 2B; Supplementary Figure 3**). Similarly, keeping a fixed amount of PTX, we titrated the amount of the GsGz chimera. We observed comparable effects with each of the four receptors indicating an optimal amount of GsGz at 45-138 ng over a total of 2.5 µg of transfected DNA (**Figure 2C; Supplementary Figure 4**). Finally, we observed that the greatest accumulation of cAMP was achieved when transfecting the highest amount of receptor (**Figure 2D; Supplementary Figure 5**). Based on these results, we recalibrated the amount of each transfected plasmid, expressing the four assay components, and determined the optimal conditions as a molar ratio of 50:47.5:1.8:0.7 (GloSensor:GPCR:GsGz:PTX) (**Figure 2E**). These optimization steps led to an overall 3-6 fold improvement in assay sensitivity measured as a fold change accumulation in cAMP. Given that HEK293 cells express Gα ^39^, we explored if transfecting cells with the catalytic subunit of OZITX instead of PTX would further boost the signal because it inhibits each G_i/o/z_ protein including G ^40^. We initially verified the effect of OZITX on the activation of wild-type G_i/o/z_ proteins and G protein chimeras using the G protein nanoBRET assay (**Supplementary Figure 6A**). Our data indicate a complete abolishment of signal initiated by wild type Gα_o_, Gα_i_, and Gα_z_, while the inhibition of the Gs-based chimeras was minimal for GsGo and GsGz, and more pronounced but incomplete for GsGi (**Supplementary Figure 6B**). We then incorporated OZITX in our assay and tested the activation of D2R, GABABR, ADRA2A, and MOR. Our findings indicate that, overall, OZITX demonstrated a comparable effect to PTX, with no significant enhancement observed in the assay. (**Supplementary Figure 6C**).

**Figure 2.**
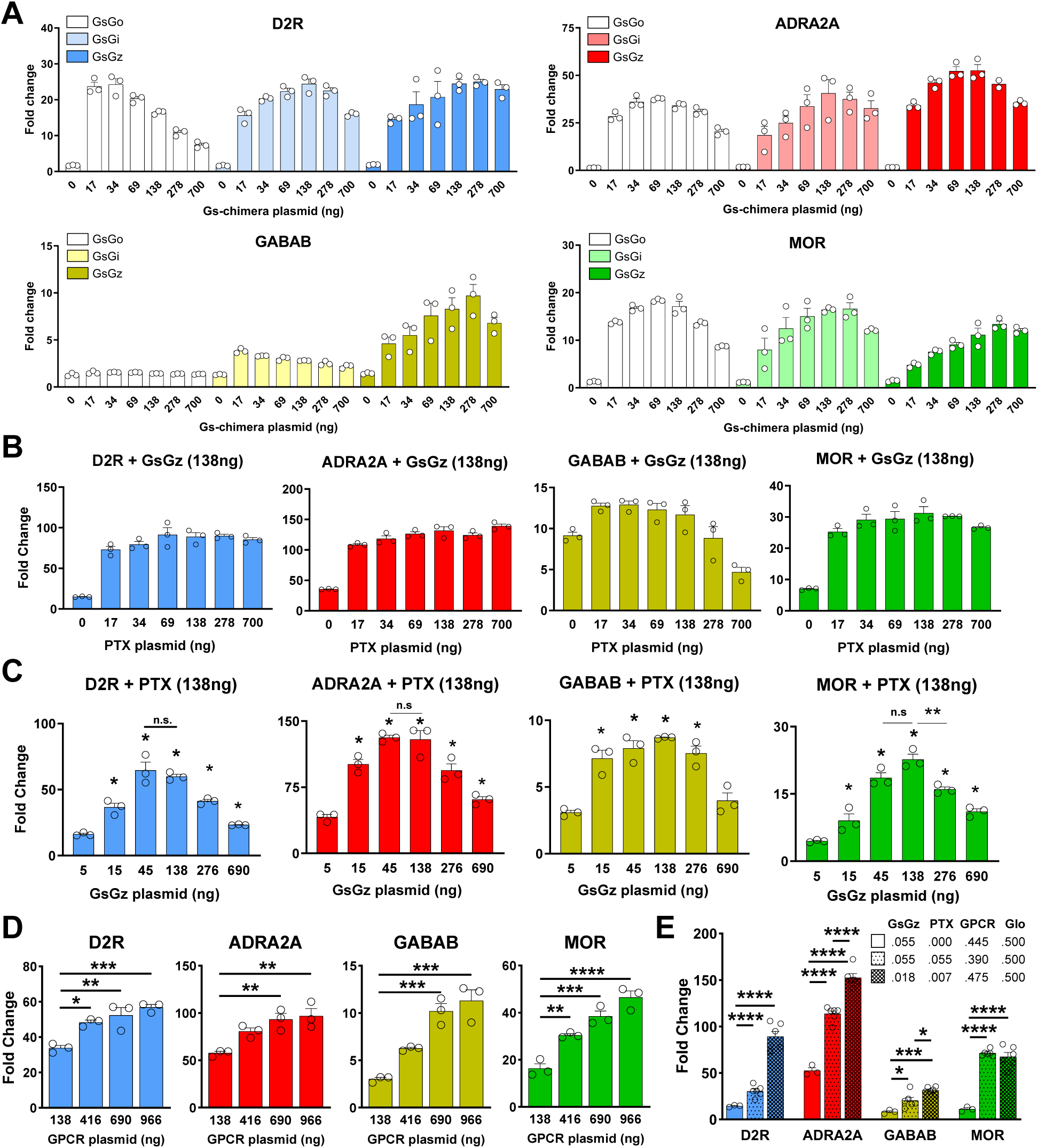
Optimization of transfection ratios between G_z_ESTY components. (**A**) Activation of indicated GPCRs with increasing amount of transfected Gs-based chimeras. (**B**) GPCR activation with the introduction of increasing concentrations of PTX-S1 subunit to eliminate the inhibition of adenylyl cyclase by endogenous G_i/o_ proteins. (**C**) GPCR activation in the presence of increasing amount of transfected GsGz plasmid. (**D**) Signal fold change with titration of each transfected receptor. (**E**) Fold-change comparison between standard transfection ratio and optimized ratio 50:47.5:1.8:0.7 (GloSensor:GPCR:GsGz:PTX). The data shown represent the average of 3-6 independent experiments (one-way ANOVA with Dunnett’s multiple comparisons test, *p < .05; **p < .01; ***p < .001; ****p < .0001). Data are shown as means ± SEM.

### 3.3 Multiple conditions can further affect signal detection in G_z_ESTY

Further conditions can significantly affect assay sensitivity. As previously reported^28^, we first confirmed that GloSensor assays produce a greater signal when performed at 28°C compared to 23°C or 37°C (**Figure 3A; Supplementary Figure 7**). Moreover, the presence of ligands in the culturing media can affect GPCR activation because of desensitization processes. We tested this factor by serum starving the cells for 4 hours or changing the cell culture media after transfection to one containing 10% charcoal-stripped FBS or 10% dialyzed FBS (**Figure 3B; Supplementary Figure 8**). Our data showed that these four prototypical GPCRs are generally not affected by the presence of normal serum, or they are negatively impacted by its absence. Furthermore, many GPCRs have been shown to interact with extracellular matrix components^41^ and, therefore, the analysis of their signaling properties may be different if studied using cells in adhesion or in suspension. We tested the same four GPCRs in these two conditions and we found that, indeed, GABAB receptor shows a significantly higher signal when cells are cultured in poly-D-lysine coated wells (**Figure 3C; Supplementary Figure 9**). Finally, we tested the possible advantage of blocking endogenous phosphodiesterases (PDEs) with IBMX to enhance the accumulation of cAMP (**Figure 3D; Supplementary Figure 10**). As expected, we found that applying IBMX before recording the baseline significantly increased the baseline reading and reduced the overall fold change observed. However, when IBMX was applied at the same time as the agonist, we measured a significant increase in signal (**Figure 3D; Supplementary Figure 10**). By subtracting the signal obtained by applying IBMX with the vehicle from the total luminesce observed, we found that the increased signal was not simply due to the simultaneous activation of adenylyl cyclase and inhibition of PDEs.

**Figure 3.**
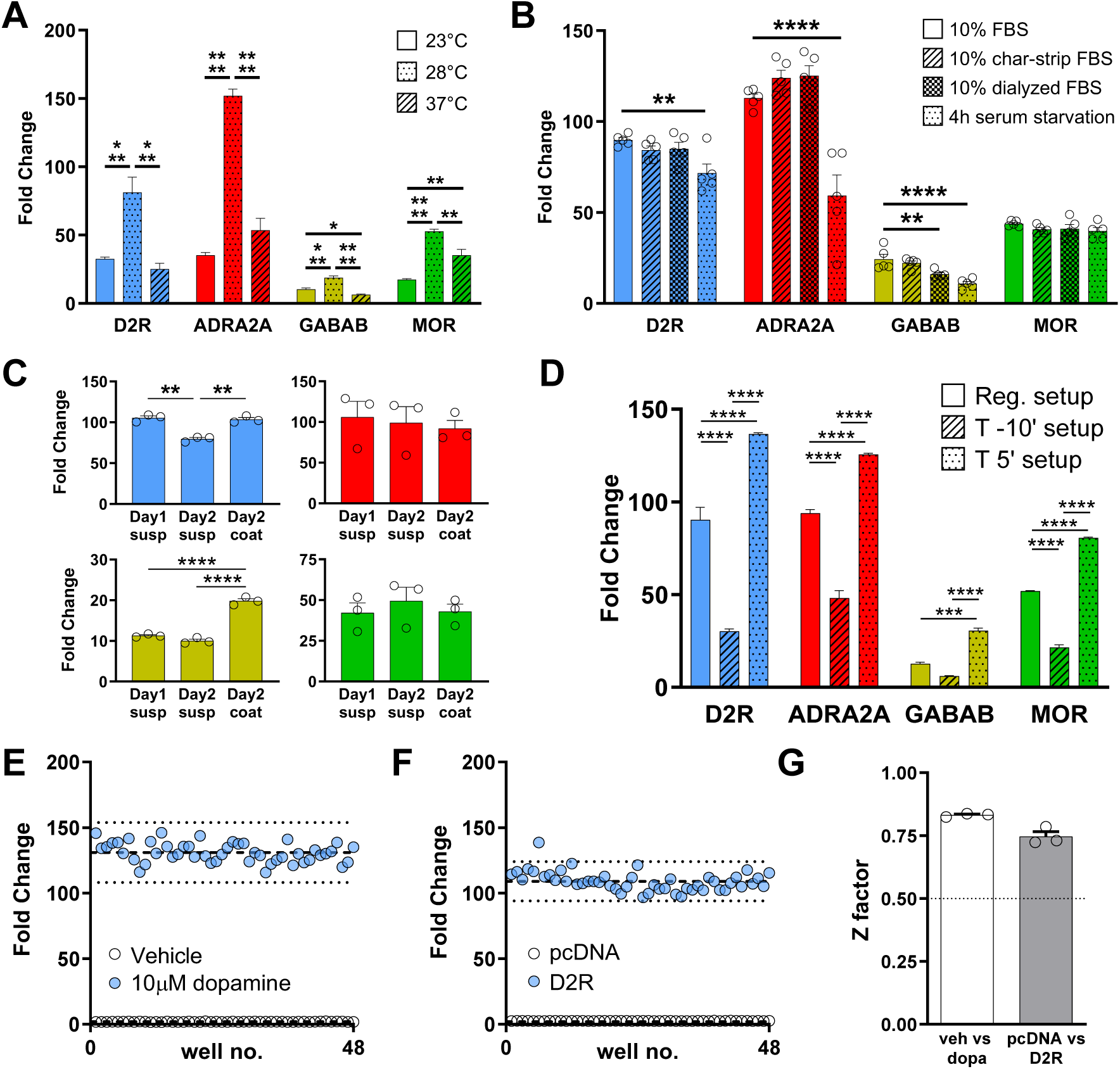
G_z_ESTY protocol optimization and Z factor calculation. (**A**) Assay temperature influences assay sensitivity. Assays performed at 28°C give a significantly higher signal. (**B**) FBS removal (4h serum starvation), use of charcoal-stripped FBS, or use of dialyzed FBS did not improve the assay signal. (**C**) Assays performed with cells in adhesion show a signal improvement for GABAB receptor but not for D2R, ADRA2A, and MOR. (**D**) Co-treatment with IBMX (T 5’ setup) significantly increases the signal with each of the four GPCRs. Data shown represent the average of 3-5 independent experiments (one-way ANOVA with Dunnett’s multiple comparisons test, *p < .05; **p < .01; ***p < .001; ****p < .0001). Data are shown as means ± SEM. (**E**) Cells transiently transfected with D2R were treated with either 10 µM dopamine or vehicle. Dashed lines represent the means of the fold change. Dotted lines display three standard deviations from the mean of each data set. (**F**) Cells transfected with indicated plasmids were treated with 10 µM dopamine. Dashed lines represent the means of the fold change for cells expressing D2R or cells not expressing exogenous GPCRs (pcDNA, control). (**G**) Z factor calculation. Dotted lines indicate the threshold for robust assays. Data shown in panels **E** and **F** are representative of three independent experiments quantified in panel **G** as means ± SEM.

To quantify the robustness of G_z_ESTY, we calculated a screening window coefficient (Z factor). This factor reflects both the dynamic range of the assay and its variability, and it is a significant metric of the assay quality. Considering the overall goal of using G_z_ESTY for GPCR deorphanization where agonists are normally not available, we calculated the Z factor in two different conditions. We first compared cells transfected with D2R or GABAB receptors and treated with the respective agonist vs vehicle (**Figure 3E; Supplementary Figure 11A**). Then, we analyzed the effect of agonist treatments on cells expressing each GPCR vs mock-transfected cells (**Figure 3F; Supplementary Figure 11B**). This condition mimics the conditions for high-throughput screening to identify orphan GPCR ligands. Independent replicates conducted on different days demonstrated that the assay is highly reproducible, with Z factors calculated at 0.75 and 0.83 for D2R, and 0.78 for both conditions for the GABAB receptor (**Figure 3G; Supplementary Figure 11C**).

### 3.4 Generation of an all-in-one plasmid for G_z_ESTY cell delivery

Like most cell-based systems used to detect GPCR signaling, G_z_ESTY is a multicomponent assay (**Figure 4A**). As a result, functional measurements require efficient co-delivery of multiple plasmids in each cell. This issue is normally overcome by averaging the signal of a large number of cells assuming that a significant fraction will express all the components. We recently generated a facile system to produce large plasmids encoding multiple proteins under the control of individual promoters. This approach guarantees that each transfected cell will express all of the necessary components, thereby increasing the assay’s sensitivity and simplifying transfection protocols. To select promoters to drive the expression of each component recapitulating our previously defined ratios, we performed a promoter efficiency assay (**Figure 4B**). Based on these results, we selected a minimal CMV promoter to express the GloSensor and GPCRs, and a UbC promoter for the expression of PTX-S1 and GsGz. We subcloned each component accordingly and generated a set of all-in-one plasmids (**Figure 4C**). We then tested these newly generated tools in cells transiently transfected with each of our four prototypical G_i/o/z_-coupled receptors that we stimulated with selective agonists. We found that the 4-in-one plasmids performance was comparable to the co-transfection of multiple plasmids, but greatly simplified transfection protocols and reproducibility (**Figure 4D**).

**Figure 4.**
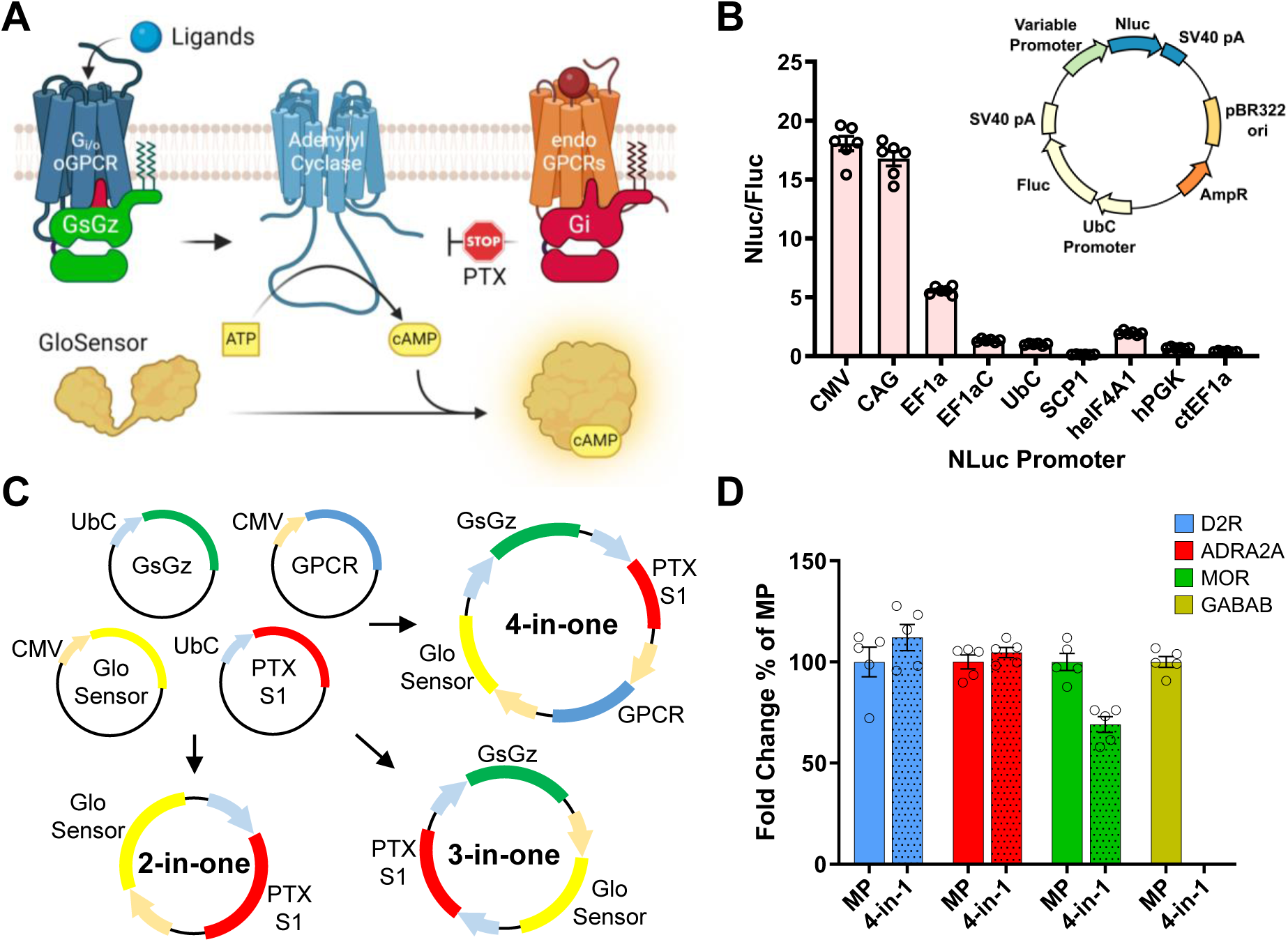
All-in-one G_z_ESTY plasmids. (**A**) Schematics of G_z_ESTY and each of its components. (**B**) Promoter efficiency estimation calculated as Nluc luminescence normalized over firefly luminescence signal in HEK293 cells transfected with a plasmid encoding nanoluc under the control of each of the indicated promoters and firefly under the control of UbC promoter. N=6 (**C**) Maps of the all-in-one plasmids generated. (**D**) Assay comparison in cells transiently transfected with four single plasmids or with each of the 4-in-one plasmids encoding each indicated GPCR. Data are shown as means ± SEM; N=5.

Next, we expanded the analysis of G_z_ESTY capabilities to a larger array of 24 G_i/o/z_-coupled GPCRs paired with the 3-in-one construct. We confirmed the suitability of the assay to study the pharmacology of 23 out of 24 G_i/o/z_-coupled receptors (**Figure 5A** and **5C**). At the same time, we confirmed that in our setup we can still detect the activation of 5 G_s_-coupled receptors (**Figure 5A**). Overall, this data demonstrated that G_z_ESTY is a powerful tool for GPCR deorphanization as it sensitively detects activation of both G_i/o/z_- and G_s_-coupled GPCRs, potentially enabling the study of 86% of the GPCRome and, by similarity, most of the orphan GPCRs (**Figure 5B**). We next investigated if G_z_ESTY was suitable to detect partial agonism. Using MOR as a model, we obtained concentration-response curves after stimulation with the endogenous ligand β-endorphin (pEC50=5.45, E_max_=62.91), with the full agonist DAMGO (pEC50=7.17; E_max_=59.53), and with the partial agonist morphine (pEC50=7.21, E_max_=38.85) that showed 65% of the maximal amplitude measured with DAMGO (**Figure 6A**). We also demonstrated that G_z_ESTY can be used to study the activation of endogenously expressed receptors. To this goal, concentration-response curves were obtained from cells transfected with the 3-in-one construct, which does not express any exogenous GPCR, and treated with selective peptidic agonists activating endogenously expressed protease-activated receptor (PAR) PAR1 (pEC50=4.56, E_max_=73.87) or PAR2 (pEC50=4.97, E_max_=103.70). Cells transfected with the 2-in-one construct (no GsGz chimera) served as negative control (**Figure 6B**).

**Figure 5.**
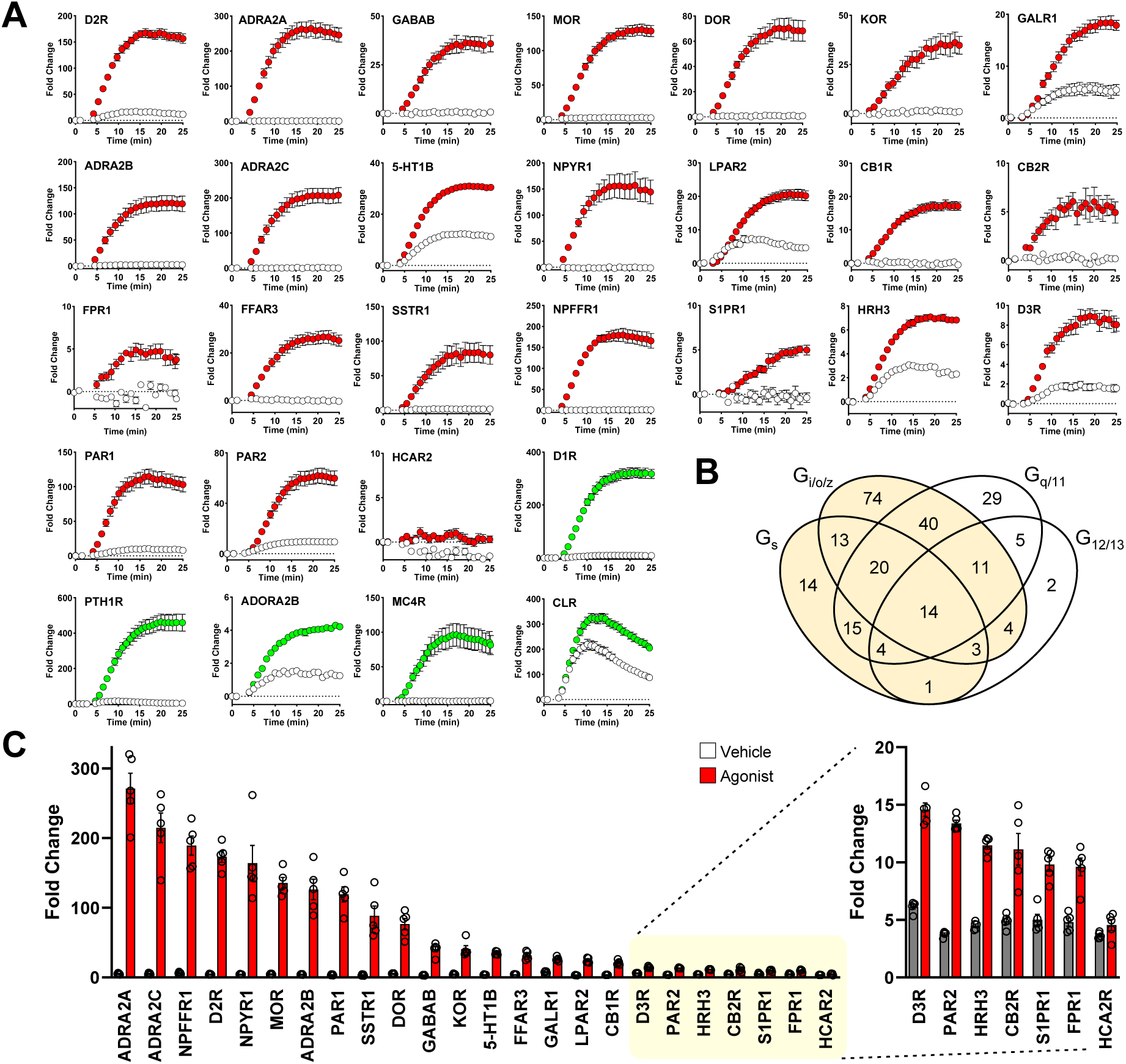
G_z_ESTY applied to a battery of ligand-activated GPCRs. (**A**) Real-time analysis of ligand-mediated activation of a battery of G_i/o/z_-PCRs (red) and G_s_-PCR (green) in cells co-transfected with indicated receptors and the 3-in-one G_z_ESTY plasmid in the presence of 50 µM IBMX. Cells transfected only with the 3-in-one plasmid served as control (white). The following agonists were applied at 4 minutes: 10 µM dopamine (D2R, D1R), 100 µM clonidine (ADRA2A, ADRA2B, ADRA2C), 10 µM GABA (GABABR), 10 µM DAMGO (MOR), 1 µM SNC-80 (DOR), 10 µM salvinorin A (KOR), 10 µM human galanin (1-30) (GAL1R), 100 µM serotonin (5-HT1B), 10 µM human neuropeptide Y (13-36) (NPYR1), 10 µM lysophosphatidic acid (LPAR2), 10 µM 2-arachidonoyl glycerol (CB1R, CB2R), 10 µM N-formyl-met-leu-phe (FPR1), 1 mM isobutyric acid (FFAR3), 10 µM somatostatin 14 (SSTR1), 10 µM neuropeptide FF (NPFFR1), 10 µM SEW2871 (S1PR1), 10 µM histamine (HRH3), 10 µM quinpirole (D3R), 10 µM TFLLR (PAR1), 10 µM SLIGKV (PAR2), 10 µM MK-6892 (HCA2R), 1 µM teriparatide PTH (PTH1R), 10 µM AB-MECA (ADORA2B), 10 µM NDP-α-MSH (MC4R), and 10 µM CGRP (CLR). Data are shown as means ± SEM; N=5. (**B**) GPCRs that couple to G_s_ and/or G_i/o/z_ and can potentially be detected by GZESTY are highlighted and include 213 out of 249 total ligand-activated GPCRs (86%) (adapted from GPRCdb.org). (**C**) Quantification of agonist-induced activity for 24 G_i/o/z_-coupled GPCRs. On the right, the response to agonist of the last seven GPCRs is also reported with a different scale. Data are shown as means ± SEM of the fold change obtained; N=5.

**Figure 6.**
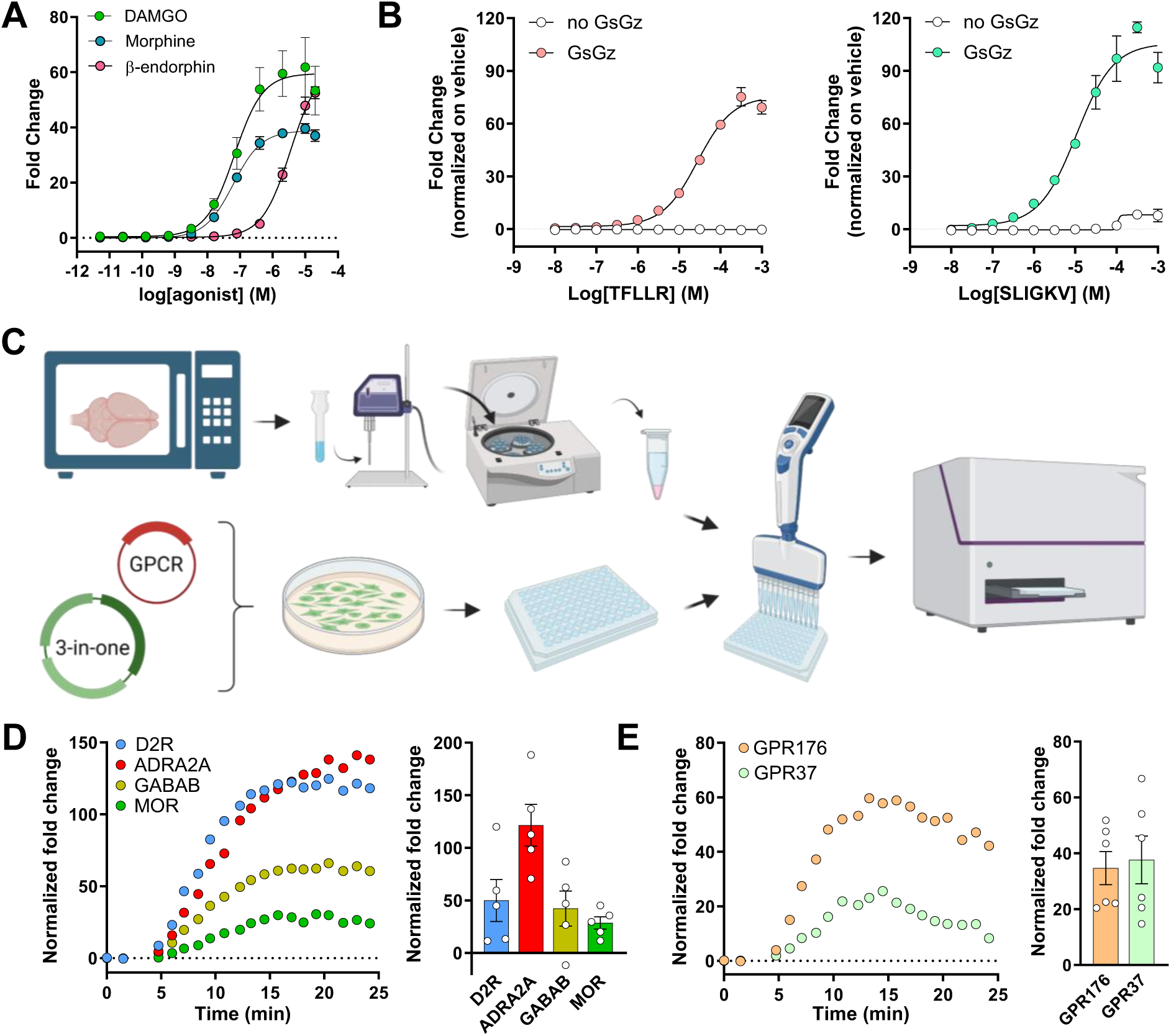
G_z_ESTY applications. (**A**) Concentration-response curves obtained for MOR stimulated with DAMGO, morphine, or β-endorphin. Data are shown as means ± SEM; N=3. (**B**) Concentration-response curves of endogenously expressed PAR1 activated with a selective agonist peptide TFLLR (left) or PAR2 activated with selective agonist peptide SLIGKV (right) in cells transfected only with the 3-in-one plasmid. Control cells were transfected with a 2-in-one plasmid not expressing GsGz chimera. Data are shown as means ± SEM; N=3. (**C**) Schematics of G_z_ESTY application to test ligand presence in mouse brain extract. Mouse brains were isolated after 10’’ of microwave exposure to preserve peptides and small molecules from degradation. Brain homogenates were sonicated, and debris and insoluble materials were separated by centrifugation. Supernatant was then applied to cells transfected with G_z_ESTY and resuspended in 96-well plates. cAMP levels were measured before and after brain extract application for a total of 25 minutes (created with BioRender). (**D**) Crude brain extract application induces activation of D2R, ADRA2A, MOR, and GABAB receptors. Representative traces (left) and quantification of 5 independent experiments (right) indicate the presence of endogenous ligands in the brain extract. Data are shown as means ± SEM; N=5. (**E**) Representative traces (left) and quantification (right) indicating the activation of GPR176 and GPR37 by crude brain extract application. Data are shown as means ± SEM; N=6.

### 3.5 G_Z_ESTY application to GPCR deorphanization

To assess the suitability of G_z_ESTY in projects aiming at the identification of endogenous ligands for orphan GPCRs, we treated cells co-transfected with the 3-in-one plasmid and individual GPCRs with a raw source of endogenous ligands. Historically, similar strategies were successfully used to isolate anandamide from pig brain, identifying it as an endogenous ligand for cannabinoid receptors^42^; these methods also facilitated the identification of ghrelin from rat stomach extract^43^, and helped pair other GPCRs with their endogenous ligands^44–47^. To this goal, we prepared a crude aqueous mouse brain extract that we applied to G_z_ESTY-transfected cells, and we measured cAMP accumulation over time (**Figure 6C**). Control cells transfected with the 3-in-one plasmid and therefore expressing GloSensor, GsGz, and PTX, but not exogenous GPCRs, showed significant induction of cAMP levels likely due to the activation of endogenous receptors. This baseline was used to normalize the signal obtained from cells overexpressing exogenous GPCRs. Applying this strategy, we demonstrated that the high sensitivity of G_z_ESTY enables the detection of robust and consistent activation of D2R, ADRA2A, GABAB, and MOR from crude brain extracts, which likely contain highly diluted endogenous agonists (**Figure 6D**). We next apply the same strategy to explore the possible activation of a battery of orphan GPCRs from crude brain extracts. Intriguingly, we detected a consistent activation of orphan receptors GPR176 and GPR37 suggesting a significant presence of their endogenous ligands in the mouse brain (**Figure 6E**). Both these orphan GPCRs are expressed in the brain, and they are reported to couple to heterotrimeric G proteins of the G_i/o/z_ family, but their endogenous ligands have not been identified or agreed upon^48–50^. We performed parallel experiments using the same strategy to test the presence of GPCR ligands in bovine pituitary extracts. We reasoned that many GPCRs are activated by hormones released by glands, and we used the follicle-stimulating hormone (FSH) receptor as a positive control, as its endogenous ligand is secreted by the pituitary gland. We found that pituitary extract can indeed activate FSHR, and we measured some level of activation of GABAB receptor, but none of the orphan GPCRs we tested were activated (**Supplementary Figure 12**).

## 4. DISCUSSION

Orphan GPCRs, by definition, have unknown or poorly understood ligands and signaling mechanisms. Without a known ligand or baseline receptor activity, designing assays to measure their activation or inhibition becomes a complex task. A major obstacle in identifying endogenous ligands for orphan GPCRs is the absence of sensitive assays capable of detecting low concentrations of these ligands in biological samples. This challenge is especially pronounced for orphan GPCRs that couple to heterotrimeric G proteins of the G_i/o/z_ family, which, according to the G protein database ((GPCRdb.org), represent 72% of deorphanized GPCRs, with 30% being uniquely coupled to G_i/o/z_ proteins^51,52^. To address this problem, we designed G_z_ESTY, an optimized cell-based assay that enables robust detection of GPCR activation in response to both purified ligands and mixed sources of molecules, including those containing diluted ligands.

Our work identified optimal assay conditions that apply to each GPCR tested, such as the temperature of the assay, ratio of assay components, and presence of IBMX, while other parameters appear to affect the readout in a GPCR-specific manner. For example, we did not observe a significant effect of serum on the assay outcome for four prototypical GPCRs tested, however, the presence of ligands in the culture media has been shown to induce receptor desensitization and internalization halting the access of extracellular ligands to the receptor itself^53–55^. Therefore, we cannot exclude the possibility that testing certain GPCRs may benefit from using reduced serum content or dialyzed serum. This aspect will require specific assessment for individual receptors. Using IBMX has the advantage of enhancing the detected signal, thereby improving the assay’s sensitivity. Additionally, it helps eliminate false positives in high-throughput screens caused by the presence of phosphodiesterase inhibitors in many compound libraries. By applying innovative cloning techniques, we generated large plasmids that encode multiple proteins under the control of unique promoters. These all-in-one plasmids have the advantage of expressing each component at precisely defined and optimized ratios. Moreover, their use greatly simplifies transfection protocols for large-scale screening efforts. Using all-in-one plasmids, we showed that G_z_ESTY is suitable to detect the activation of endogenously expressed receptors. Delivery of the 3-in-one plasmid to relevant cell types can be combined with agonist treatment to perform pharmacological studies of receptors of interest in a physiologically relevant context. Our initial optimization steps for G_z_ESTY were performed using four prototypical GPCRs (D2R, ADRA2A, GABAB, and MOR). The optimized protocol was later applied to an array of 24 G_i/o/z_-coupled GPCRs. The results showed that the great majority exhibited over a 20-fold change in signal compared to baseline, suggesting ideal signal-to-noise properties for the assay. We also observed smaller signal amplitude for a few receptors. This lower sensitivity may depend on several factors including the presence of endogenous agonists in the culture media that can lead to receptor desensitization or the requirement of performing the assay with cells in adhesion. Transfected receptors may also not be properly expressed, targeted to the plasma membrane, or require accessory proteins to generate functional receptor complexes. Therefore, when working with a specific receptor it is recommended to test multiple assay conditions in the cell type of choice to achieve an optimal assay sensitivity.

Screening efforts to identify orphan GPCR agonists will be markedly enhanced by utilizing G_z_ESTY. Analysis of the constitutive activity of orphan GPCRs provides valuable information regarding their G protein coupling profile^35,56,57^. Recent studies have shown that, similar to ligand-activated GPCRs, the majority of orphan GPCRs couple to heterotrimeric G proteins of the G_i/o/z_ family. This makes them suitable candidates for analysis using G_z_ESTY. Furthermore, given the limited understanding of the G protein coupling profile for many orphan GPCRs, a key benefit of G_z_ESTY is its ability to detect the activation of approximately 86% of GPCRs. This estimate is based on coupling data regarding the number of GPCRs that can couple to G_s_ and/or G_i/o/z_ proteins (**Figure 5B**).

The isolation of orphan GPCR endogenous ligands relies significantly on the availability of methods that can accurately detect their presence in raw unfractionated extracts. In this study, we utilized G_z_ESTY to identify the existence of endogenous ligands for orphan GPCRs in the mouse brain. Our approach consistently detected known neurotransmitters capable of activating adrenergic, dopaminergic, GABAergic, and opioid receptors, thereby validating the method. Notably, we also identified the presence of ligands that can activate class A orphan receptors GPR176 and GPR37. GPR176 was previously found to be enriched in the mouse suprachiasmatic nucleus (SCN) of the hypothalamus, the central pacemaker that controls the circadian rhythm^48,58^. Its protein expression levels oscillate according to the circadian period being higher during the night, and GPR176-deficient mice display a significantly shorter circadian period compared to their wild-type littermates. Further studies indicated that GPR176 is probably not a light signal-related receptor for the SCN; rather, it seems to be involved in determining the intrinsic period of the SCN. Finally, heterologous expression of GPR176 blunted the forskolin-induced accumulation of cAMP suggesting its coupling to heterotrimeric G_i/o/z_ proteins^48^. Overall, these findings indicate that GPR176 is involved in the suppression of cAMP production in the SCN during nighttime, which serves as the molecular mechanism for its regulation of the circadian period^59^. GPR176 roles outside the central nervous system have also been described^60^; however, no endogenous or synthetic ligands capable of activating GPR176 have been identified yet. With G_Z_ESTY enabling the detection of GPR176 endogenous ligand in the mouse brain, we will be able to perform further studies leading to its isolation. GPR37 is a brain-enriched orphan GPCR with some of the highest transcript levels among all GPCRs^61^. Despite its well-studied involvement in Parkinson’s disease, inflammatory responses, and pain, the identity of its endogenous ligand remains a topic of debate^49,62–64^. Early studies proposed prosaptide, a fragment of the secreted neuroprotective and glioprotective factor prosaposin, as the endogenous ligand for GPR37^50^. The identity of this ligand-receptor pair was supported by the fact that prosaptide stimulation of cells transfected with GPR37 induced ERK phosphorylation in a PTX-sensitive manner suggesting coupling to G_i/o_ proteins^50^. Moreover, ^35^S-GTPγS binding assays and inhibition of forskolin-induced cAMP accumulation all pointed at prosaptide as the endogenous ligand for GPR37^50^. GPR37 activation by prosaptide was later confirmed by measuring a PTX-sensitive intracellular calcium mobilization in GPR37 transfected HEK293 cells^65^. In the same study, activation by neuroprotectin D1 was also observed. Studies in astrocytes also detected GPR37 activation by prosaptide^66^. However, the claim that prosaposin and prosaptide are the endogenous ligands for GPR37 was later challenged by reports demonstrating that GPR37 shows high constitutive activity and that prosaptide treatments were ineffective (Smith, 2015). Consequently, the prosaptide/GPR37 pairing has yet to be approved by the International Union of Basic and Clinical Pharmacology (IUPHAR) Nomenclature Committee. Given that multiple endogenous ligands can activate the same receptor^8^, the pursuit of identifying GPR37 ligands detected using G_Z_ESTY is a valuable endeavor.

## Supporting information

GzESTY supplemental figures

## Author Contributions

Experimental investigation and data analysis, L.F., J.J.P., J.D.L. and C.O; Conceptualization, L.F. and C.O.; writing and editing—original draft preparation, C.O.; All authors have read and agreed to the published version of the manuscript.

