## Supplementary material for "Gz Enhanced Signal Transduction assaY (G_Z_ESTY) for GPCR deorphanization": GzESTY supplemental figures

#### Supplementary Figure 1

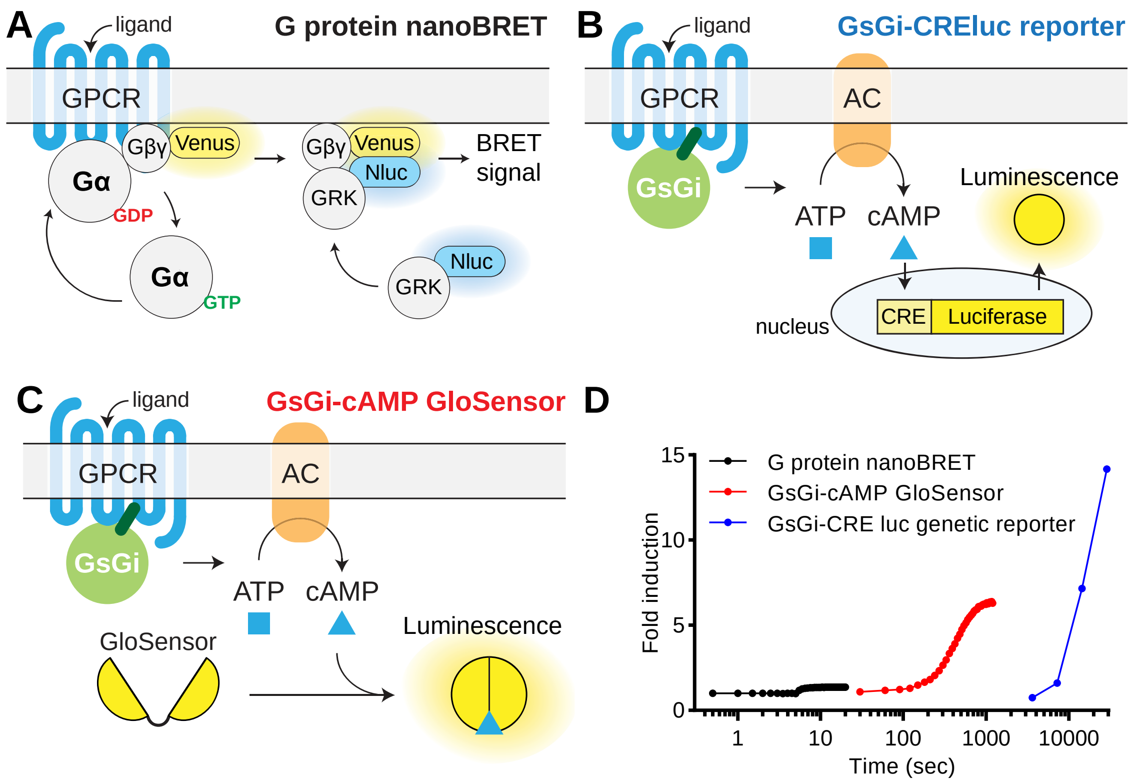


**Supplementary Figure 1. Cell-based assays to study the activation of $G_{i/o/z}$-coupled receptors.** (**A**) Schematics of the G protein nanoBRET assay. Ligand interaction with the GPCR induces the dissociation of Gβγ-Venus from Gα. Free Gβγ-Venus interacts with a plasma membrane-anchored biosensor made of the C-terminus of GRK3 fused to nanoluc. An increase in BRET signal is recorded in response to GPCR activation. (**B**) A $G_{i/o/z}$-coupled GPCR is transfected with a Gs-based chimera bearing the C-terminus of $G_{i/o/z}$ family members and is co-transfected with a CRE-dependent inducible luciferase reporter. Agonist activation of the receptor induces a concentration-dependent intracellular accumulation of luciferase. (**C**) Second messenger (cAMP) accumulation is measured using a genetically-encoded GloSensor in response to agonist treatment in the presence of a GsGi chimera. (**D**) Timescale and magnitude (fold induction over basal signal) of signal detection for the described cell-based assays after GABAB receptor activation with 10 µM GABA. G protein nanoBRET (black); GsGi-cAMP GloSensor (red); and GsGi-CREluc reporter (blue).

### Supplementary Figure 2

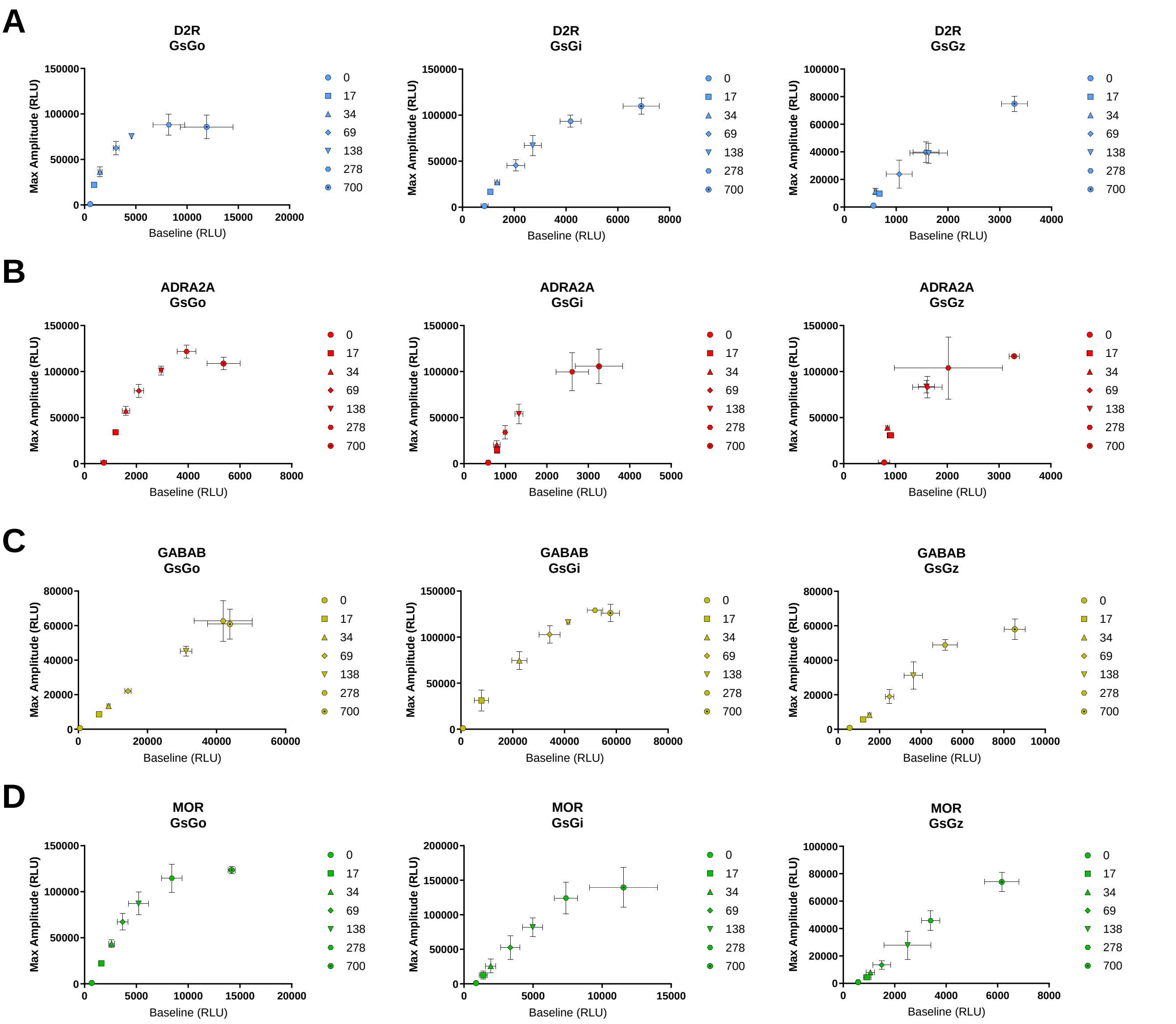


**Supplementary Figure 2.** Relationship between baseline and maximal amplitude measured as cAMP accumulation in response to receptor stimulation in the presence of increasing amount of transfected G protein chimeras (GsGo, GsGi, and GsGz). Data for D2R activated with 10 µM dopamine (**A**), ADRA2A with 10 µM clonidine (**B**), GABABR with 10 µM GABA (**C**), and MOR with 1 µM DAMGO (**D**). N=3 independent replicates.

### Supplementary Figure 3

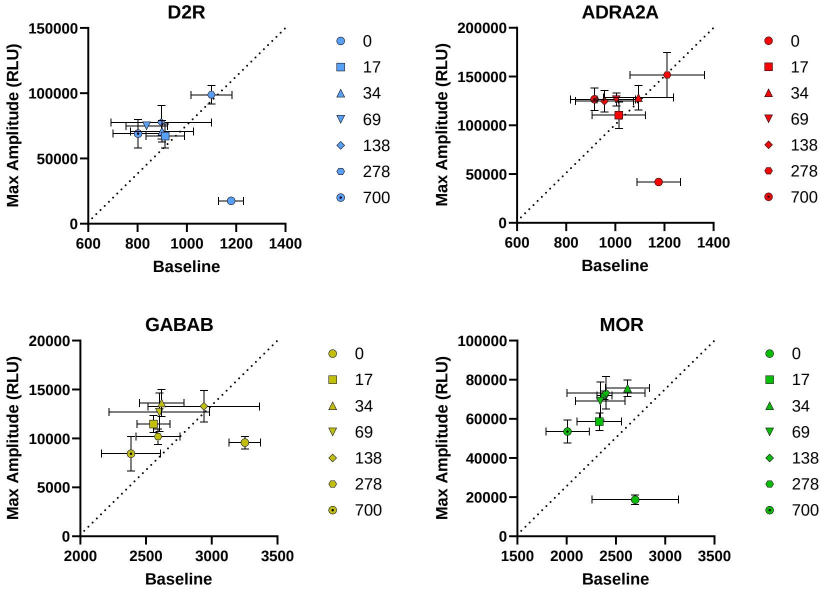


**Supplementary Figure 3.** Correlation between baseline and maximal amplitude measured as cAMP accumulation in response to stimulation of indicated receptors in cells transfected with indicated GPCRs (416 ng), GsGz chimera (138 ng), and increasing amounts of plasmid expressing PTX-S1 as reported. N=3 independent replicates.

### Supplementary Figure 4

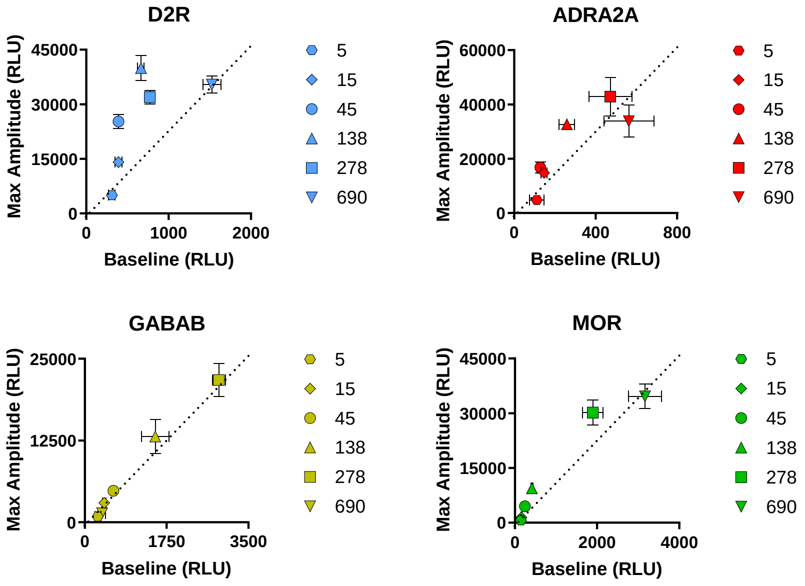


**Supplementary Figure 4.** Relationship between baseline and maximal amplitude measured as cAMP accumulation in response to receptor stimulation in cells transfected with indicated GPCRs (416 ng), PTX (138 ng), and increasing amounts of plasmid expressing GsGz chimera as reported. N=3 independent replicates.

### Supplementary Figure 5

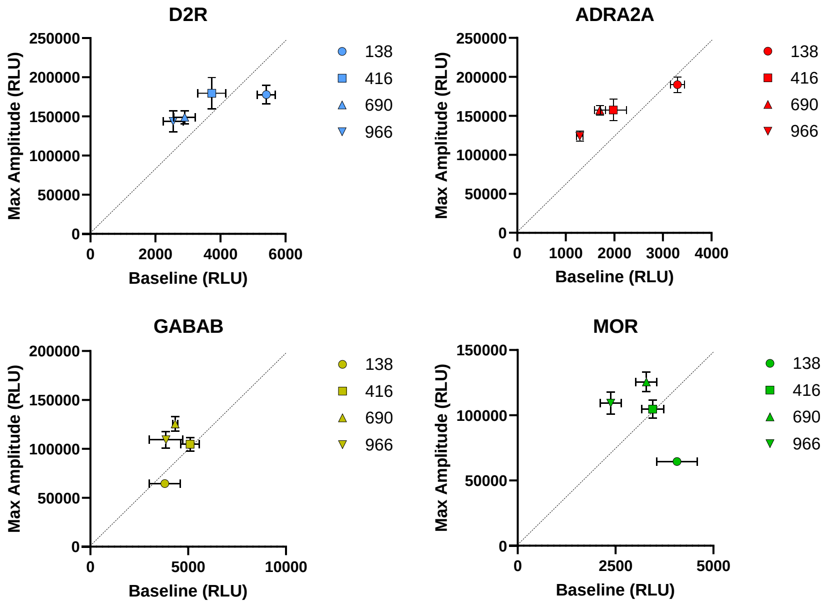


**Supplementary Figure 5.** Correlation between baseline and maximal amplitude measured as cAMP accumulation in response to stimulation of indicated receptors. Cells were transiently transfected with GsGz chimera (138 ng), PTX-S1 (138 ng), and increasing amounts of each receptor as reported. N=3 independent replicates.

### Supplementary Figure 6

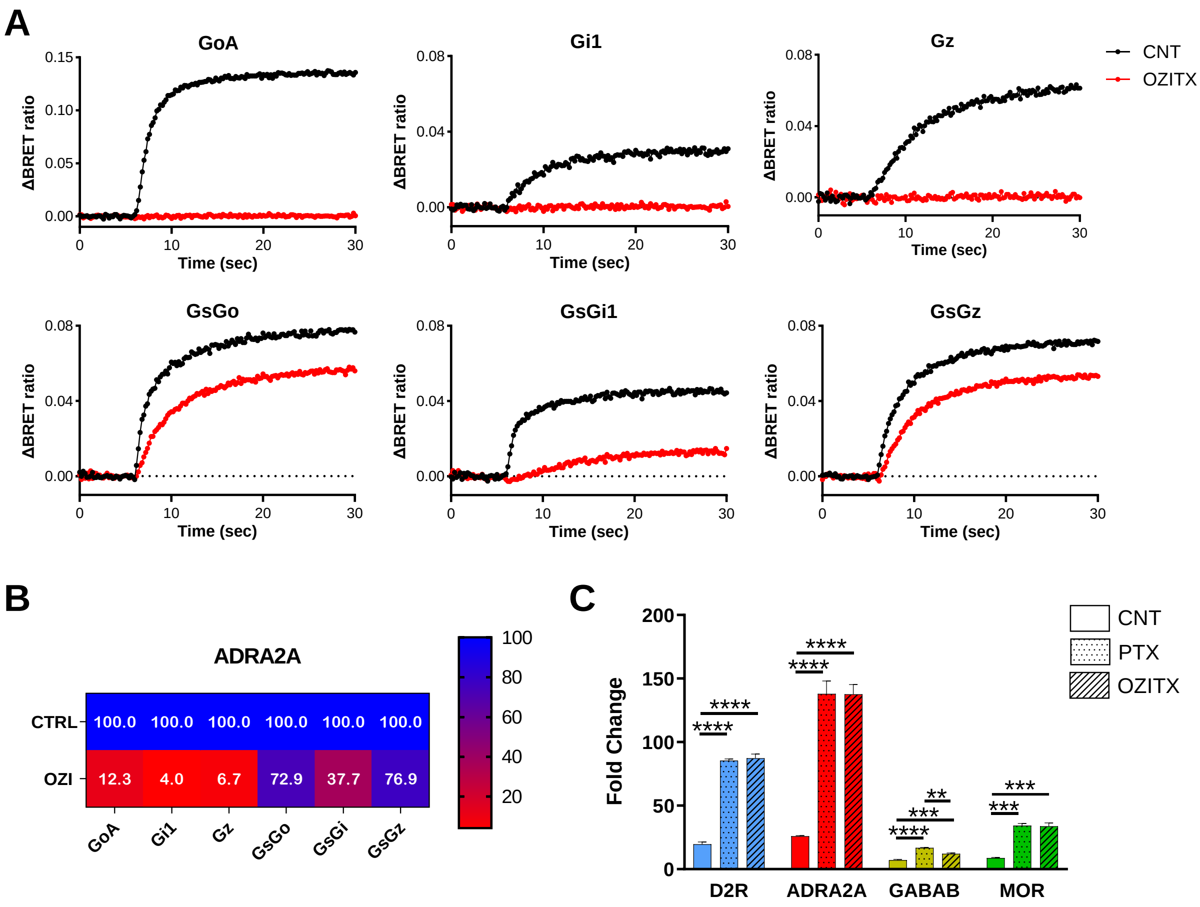


**Supplementary Figure 6.** (**A**) G protein nanoBRET analysis of the activation of wildtype G proteins (GoA, Gi1, and Gz) and G protein chimeras (GsGo, GsGi1, and GsGz) by ADRA2A in response to 10 µM clonidine with or without OZITX. (**B**) Quantification of signal inhibition by OZITX. (**C**) Fold change quantification of GzESTY performed with each of the indicated GPCRs with PTX or OZITX versus control (CNT). Data are shown as means ± SEM; N=3 (one-way ANOVA with Dunnett's multiple comparisons test, $^{**}p < .01$; $^{***}p < .001$; $^{****}p < .0001$).

#### Supplementary Figure 7

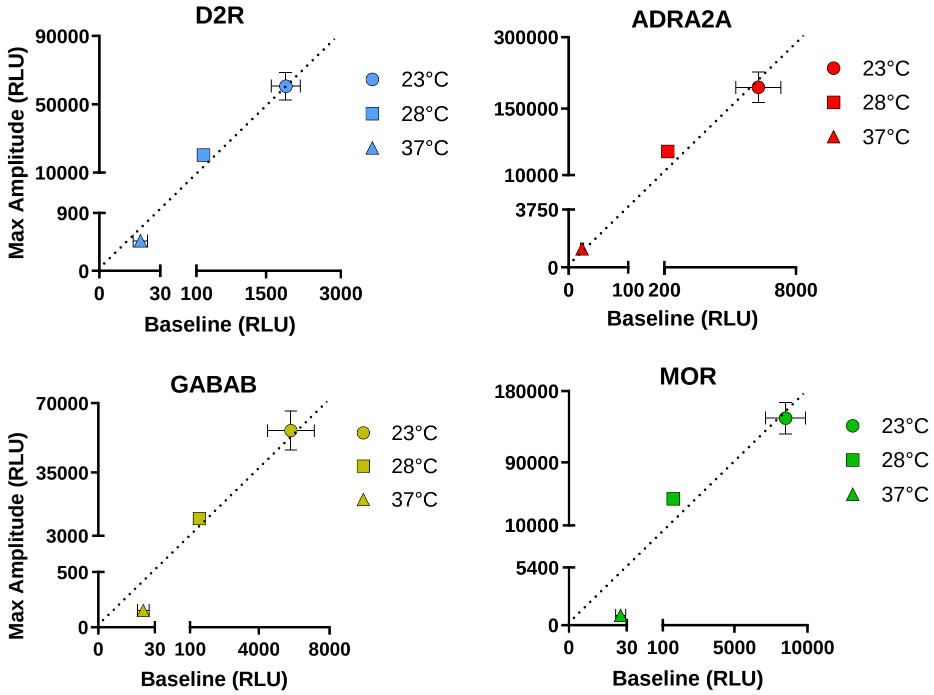


**Supplementary Figure 7.** Correlation between baseline and maximal amplitude measured as cAMP accumulation in response to stimulation of indicated receptors at 23°C, 28°C, or 37°C. Data are shown as means ± SEM; N=5.

### Supplementary Figure 8

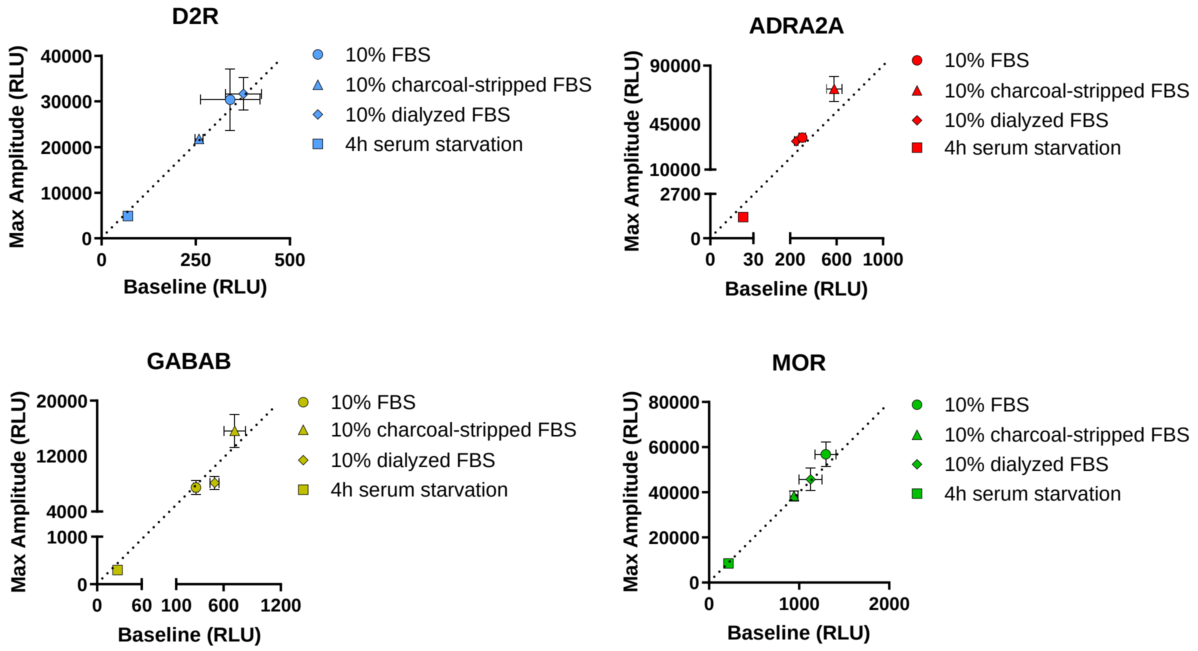


**Supplementary Figure 8.** Correlation between baseline and maximal amplitude measured as cAMP accumulation in response to stimulation of indicated receptors in transfected cells that were cultured overnight in medium containing 10% FBS (circles), 10% charcoal-stripped FBS (triangles), 10% dialyzed FBS (diamonds), or serum starved for 4 hours before functional assays (squares). Data are shown as means ± SEM; N=5.

### Supplementary Figure 9

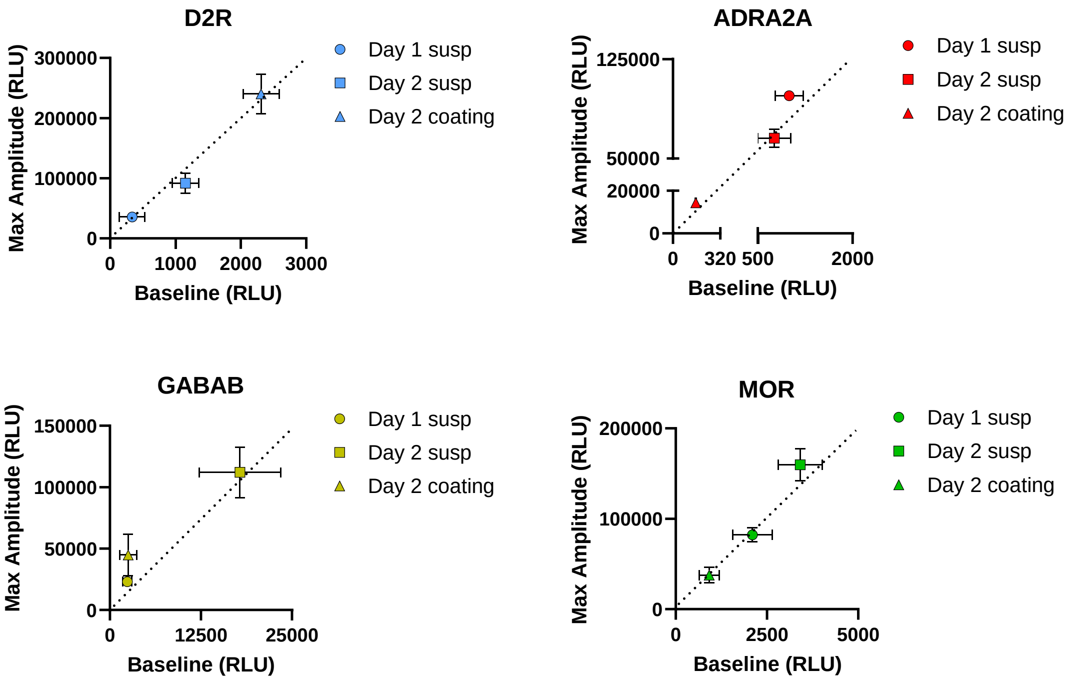


**Supplementary Figure 9.** Correlation between baseline and maximal amplitude measured as cAMP accumulation in response to stimulation of indicated receptors in transiently transfected cells. After transfection, cells were cultured for 24 hours and resuspended before the experiment (circles), cultured for 48 hours and resuspended before the experiment (squares), or cultured for 48 hours on poly-D-lysine coated wells and used without resuspension (triangles). Data are shown as means ± SEM; N=3.

### Supplementary Figure 10

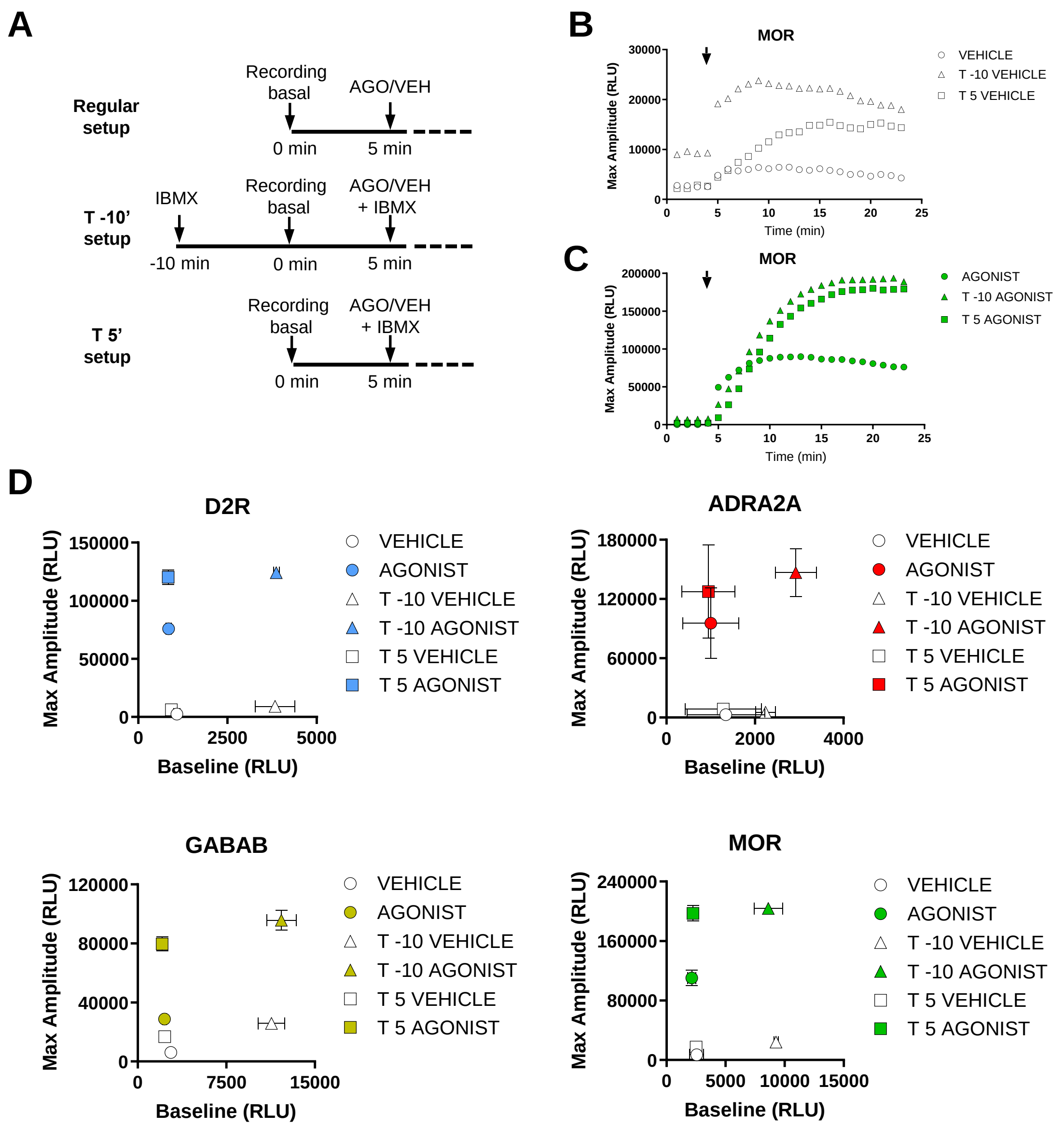


**Supplementary Figure 10.** (**A**) Assay design to test the effect of phosphodiesterase inhibition with IBMX on fold change. 50 µM IBMX was applied 10 minutes before recording the basal signal (T -10' setup) or at the same time as agonist/vehicle addition (T 5' setup). Representative traces of cAMP accumulation over time in response to MOR stimulation with vehicle (**B**) or 1 µM DAMGO (**C**) at the time indicated with an arrow. (**D**) Correlation between baseline and maximal amplitude measured as cAMP accumulation in response to stimulation of indicated receptors in transfected cells. Conditions tested included no IBMX (circles), IBMX pre-treatments for 10 minutes before agonist/vehicle stimulation (triangles), or IBMX co-treatment with agonist/vehicle (squares). Data are shown as means ± SEM; N=3.

### Supplementary Figure 11

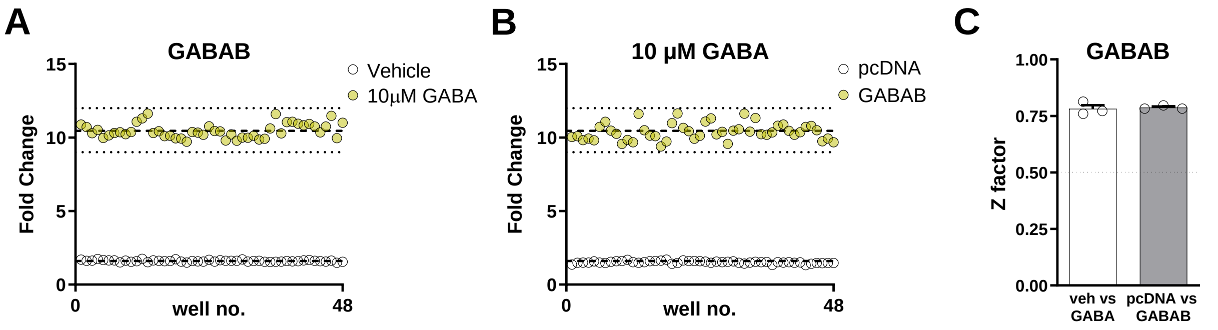


**Supplementary Figure 11. Z factor calculation for $G_Z$ESTY.** (**A**) Cells transiently transfected with GABAB were treated with 10 µM GABA or vehicle. Dashed lines represent the means of the fold change. Dotted lines display three standard deviations from the mean of each data set. (**B**) Transfected cells were treated with 10 µM GABA. Dashed lines represent the means of the fold change for cells expressing GABAB receptor or cells not expressing exogenous GPCRs (pcDNA, control). Dotted lines display three standard deviations from the mean of each data set. (**C**) Z factor calculation. Dotted lines indicate the threshold for robust assays. Data shown in panels **A** and **B** are representative of three independent experiments quantified in panel **C** as means ± SEM.

#### Supplementary Figure 12

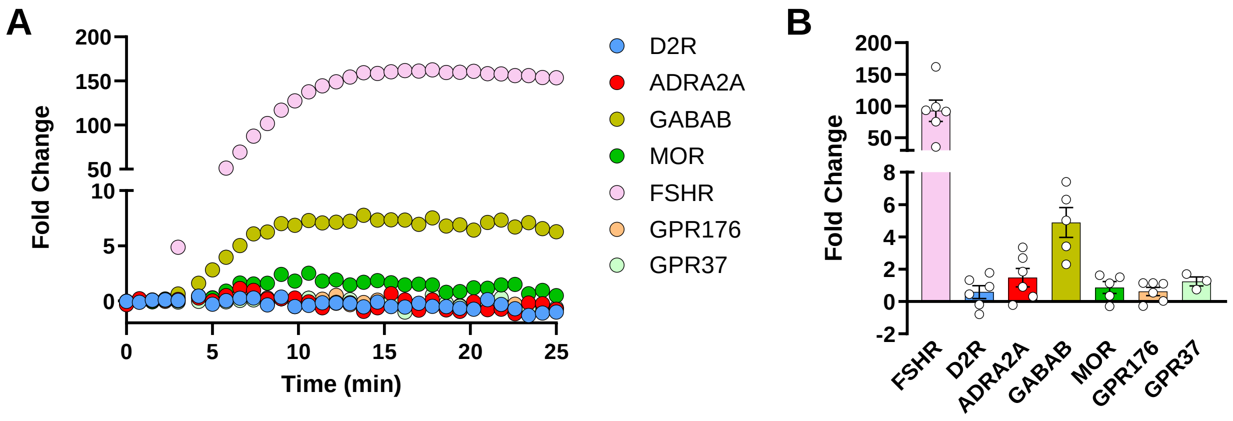


**Supplementary Figure 12.** Analysis of GPCR activation by application of bovine pituitary extract using $G_Z$ESTY. (**A**) Representative traces showing the fold change (normalized over the signal obtained in control cells not expressing exogenous GPCRs) over time in response to pituitary extract application. FSH receptor was used as a positive control. (**B**) Quantification of GPCR activation across replicates. Data are shown as means ± SEM; N=3-6.
